# Xylazine potentiates the interoceptive effects of fentanyl in male and female rats

**DOI:** 10.1101/2025.02.07.637099

**Authors:** Brooke N. Bender, Joseph M. Carew, Madigan L. Bedard, Zoe A. McElligott, Joyce Besheer

## Abstract

**Rationale:** Xylazine, a sedative typically used in veterinary medicine, has been increasingly detected as an adulterant in the unregulated opioid supply and present in opioid overdose deaths. Therefore, xylazine-adulterated fentanyl is a growing public health concern. People who use drugs have reported that xylazine changes and prolongs the effects of fentanyl

**Objectives:** We used standard operant drug discrimination procedures to better understand how xylazine impacts the discriminative stimulus/interoceptive effects of fentanyl.

**Methods:** Male and female Long-Evans rats (n=23) were trained to discriminate fentanyl (0.032 mg/kg intraperitoneal) such that one lever was reinforced with sucrose on days when fentanyl was administered, and the other lever was reinforced when vehicle was administered. Once rats met testing criteria, we tested a dose range of fentanyl to confirm discriminative stimulus control, then we tested if xylazine alone produced fentanyl-like effects and if the addition of xylazine to fentanyl impacted fentanyl interoceptive effects.

**Results:** Stimulus control was confirmed, as rats showed increased percent responses on the fentanyl-appropriate lever as well as decreased response rates for increasing doses of fentanyl. Xylazine alone did not substitute for the stimulus effects of fentanyl but produced similar response rate reductions as fentanyl alone. Xylazine co-administered with fentanyl potentiated the stimulus effects of lower doses of fentanyl in both males and females and potentiated response rate reductions.

**Conclusions:** These results indicate that xylazine enhances the interoceptive effects of fentanyl, which may inform clinical research about xylazine-adulterated fentanyl.

## Introduction

In the United States, 6 million adults and adolescents are estimated to be living with an opioid use disorder (OUD) (Keyes et al. 2022). Illicitly manufactured synthetic opioids, such as fentanyl, are relatively inexpensive and accessible (D’Orazio et al. 2023; Shafi et al. 2022). However, because people who use drugs are often unsure of the identity, potency, and potential adulterants present in the unregulated drug supply, the increased prevalence of these drugs present a major public health concern (D’Orazio et al. 2023). Xylazine is increasingly being detected as an adulterant in opioid overdose deaths in the United States, particularly along with fentanyl and most commonly in northeastern states (Friedman et al. 2022; Johnson et al. 2021; Kariisa et al. 2023; Nunez et al. 2021). In 21 jurisdictions across 20 states and the District of Columbia, the monthly percentage of deaths involving illicitly manufactured fentanyl with xylazine detected increased by over three-fold from January 2019 to June 2022 (Kariisa et al. 2023). However, because xylazine is not always included in drug tests investigating opioid overdose deaths, its prevalence across the country may be underreported (Friedman et al. 2022).

Xylazine is typically used as a veterinary anesthetic, and there are some documented instances of xylazine misuse alone (Capraro et al. 2001; Elejalde et al. 2003; Hoffmann et al. 2001; Spoerke et al. 1986). The effects of xylazine are typically attributed to its agonist activity at α-2 adrenergic receptors (Kitano et al. 2019; Schwartz and Clark 1998), but xylazine may also interact with the opioid system (Bedard et al. 2024; Browning et al. 1982; Romero et al. 2009; Sahraei et al. 2004; Samini et al. 2008). Importantly, preclinical evidence suggests that the combined sedative effects of fentanyl and xylazine, especially the induction of respiratory depression, may increase overdose risk when xylazine is administered with fentanyl (Acosta-Mares et al. 2023; Smith et al. 2023). Additionally, a recent clinical study indicated increased occurrences of recent overdose in people who reported using xylazine (Tan et al. 2024). Therefore, it is important to investigate how these drugs interact and why xylazine has emerged as a prominent adulterant in the unregulated opioid drug supply (D’Orazio et al. 2023).

Although there are several possible reasons for adulteration, adulterants are often added to enhance the effects of a drug (Cole et al. 2011). Indeed, people who use drugs and healthcare providers report that xylazine may enhance or prolong the effects of fentanyl (Friedman et al. 2022), which suggests a change in the perception of the effects of fentanyl. Interoception involves the perception and integration of bodily signals that produce an internal body state (Chen et al. 2021). Drugs elicit interoceptive effects through their physiological effects on the brain and peripheral organs, which alter bodily signals (Verdejo-Garcia et al. 2012). Different drugs produce distinct internal body states, whereas drugs with similar mechanisms of action tend to have similar interoceptive effects (Lovelock et al. 2021; Naqvi and Bechara 2010; Solinas et al. 2006; Stolerman et al. 2011). Anecdotal reports that xylazine affects the perception of fentanyl suggest xylazine may have effects that interact with the interoceptive effects of fentanyl (Friedman et al. 2022), and understanding this process as well as the receptors and brain circuitry involved has important implications on addressing the use of xylazine-adulterated fentanyl.

Operant drug discrimination procedures are well-characterized for studying the subjective/interoceptive effects of drugs (Lovelock et al. 2021; Solinas et al. 2006; Stolerman et al. 2011). These procedures train the internal drug state to serve as a discriminative stimulus to guide a specific behavior (e.g., lever selection; head entries) (Solinas et al. 2006). Moreover, drug discrimination procedures can be used to evaluate pharmacokinetics as well as the role of specific neurotransmitter systems, receptors, and brain circuitry involved in drug interoceptive effects (Besheer et al. 2003; Colpaert 1999; Jaramillo et al. 2018; Randall et al. 2020; Solinas et al. 2006). Studies employing fentanyl drug discrimination have shown that other less potent opioids administered at higher doses, such as morphine and heroin, have interoceptive effects similar to fentanyl (Colpaert and Janssen 1984; Colpaert et al. 1976a; Colpaert et al. 1975; Emmett-Oglesby et al. 1988; Flynn and France 2021; 2022; Schwienteck et al. 2019).

The effects of α-2 adrenergic receptor agonists like xylazine on fentanyl discrimination have not been evaluated. Therefore, given reports that xylazine enhances the effects of fentanyl along with evidence that it may interact with the opioid system, in the present study we investigated if xylazine alters the discriminative stimulus/interoceptive effects of fentanyl in rodents. Using standard two-lever operant drug discrimination paradigm, adult male and female rats were trained to discriminate between the interoceptive effects of fentanyl and vehicle (saline). In a series of tests, we evaluated if xylazine administration has fentanyl-like effects and if xylazine co-administration potentiates sensitivity to the interoceptive effects of fentanyl.

## Methods

### Animals

Long-Evans rats (n=24, n=12 male and n=12 female, Envigo) arrived at 7 weeks of age and were single-housed in ventilated racks with automated watering. The colony room was maintained on a 12-hour light-dark cycle, with lights on at 7am and off at 7pm, and all experiments occurred during the light cycle. Rats were given one day to acclimate to the facility, and then were handled for at least one minute for at least 5 days. Rats had *ad libitum* access to water throughout the experiment except when they were in operant chambers and *ad libitum* access to food until they reached 9 weeks of age. Three days before behavioral training, they were food restricted to 90% of their free-feeding bodyweight and fed daily at least 2 hours after training. One rat (male) was excluded from the study due to illness and was not replaced. Rats were continuously monitored and cared for by veterinary staff from the Division of Comparative Medicine at UNC-Chapel Hill. All procedures were conducted in accordance with the NIH Guide to Care and Use of Laboratory Animals and institutional guidelines.

### Drugs

All drugs were dissolved or diluted in 0.9% sterile saline and administered at 1 ml/kg via intraperitoneal (I.P.) injection. Fentanyl citrate (Spectrum Chemical) was dissolved to 0.0032, 0.005, 0.01, 0.018, and 0.032 mg/ml. Xylazine injectable solution (Covetrus, Inc.; Portland, ME) was diluted from its original concentration of 100 mg/ml to 0.032, 0.01, 0.32, and 0.1 mg/ml. When co-administered, fentanyl and xylazine were combined in a single solution at their final concentration.

### Behavioral apparatus

Experiments were conducted in 24 operant conditioning chambers (Med Associates, Georgia, VT; 31 × 32 × 24 cm) that were housed in sound-attenuating cubicles with exhaust fans and were operated using MedPC software (MedAssociates). Each chamber had bar floors, a house light on the left side, and on the right side 2 stimulus lights, each above a retractable lever, on either side of a liquid dipper receptacle housing a retractable delivery arm. Throughout the experiment, reinforcers (0.1 ml of 10% w/v sucrose solution in tap water) were delivered immediately upon the completion of the fixed-ratio (FR) schedule from a trough outside the operant chamber via the dipper arm for 4 seconds. Sucrose delivery was accompanied by illumination of the cue light above the lever. Four photobeams across the chamber were used to record beam-breaks to monitor locomotor behavior.

### Behavioral procedures

#### Lever press training

Rats first underwent 2 overnight (16-hour) lever-press training sessions, during which either the left or right lever (one each day) was inserted in the chamber, and lever presses were reinforced on increasing FR schedules (FR1 up to 70 reinforcers, FR2 up to 130 reinforcers, and FR4 for remaining reinforcers). Rats were then trained 5 days a week in 40-minute sessions, during which the house light was illuminated to indicate availability of reinforcer and a single lever (left of right, alternating daily) was inserted and reinforced on an FR schedule. Rats were trained for 4-8 days (2-4 days on each lever) on each increasing FR schedule (FR4, FR6, and FR8), and then for at least 10 days on an FR10 schedule.

#### Operant fentanyl discrimination training

After lever press training was concluded, fentanyl discrimination training began. Fentanyl discrimination was conducted similarly to our previously published alcohol discrimination studies(Besheer et al. 2009; Tyler et al. 2022). Rats were injected (I.P.) with either fentanyl or saline and were immediately placed in their assigned chamber for a 10-minute timeout period with no levers inserted. After the timeout period, both levers were inserted and the house light was illuminated, signaling the start of the 15-minute training session. Only one lever (active) was reinforced on an FR10 schedule. For each rat (counterbalanced between rats), one lever was assigned as the active lever on fentanyl days (e.g., right lever), and the other lever was assigned as the active lever on saline days (e.g., left lever).

Training sessions occurred daily Monday-Friday on a double alternating schedule such that fentanyl and saline were injected twice every four training days (e.g., saline, saline, fentanyl, fentanyl, …). The length of the timeout period (10 minutes) and the initial fentanyl training dose (0.01 mg/kg) were chosen based on previous operant fentanyl discrimination literature (Flynn and France 2021; 2022). During these sessions, the percent of responses on the fentanyl-appropriate lever for the entire session and prior to the first reinforcer and response rates were calculated. Criteria to progress from training to testing (testing criteria) was >80% of responses on the appropriate lever (>80% on the fentanyl lever on fentanyl sessions and <20% on the fentanyl lever on saline session) in the total session and prior to the first reinforcer for 8 of 10 consecutive training sessions. Because few rats (∼25%, 6 males) met testing criteria after 84 training sessions (42 each of 0.01 mg/kg fentanyl and saline), the training dose of fentanyl was increased to 0.032 mg/kg to improve discrimination performance. After 28 more training sessions, several rats (n=10, 6 female and 4 male) did not meet testing criteria due to either poor discrimination or suppression of response rates. These rats (Cohort A) continued training for 20 days on a new, intermediate training dose (0.018 mg/kg fentanyl) to evaluate its potential as a future training dose to promote adequate discrimination without suppressing behavior. The remaining rats (Cohort B) that met testing criteria after 0.032 mg/kg fentanyl discrimination training (n=13, 6 female and 7 male) proceeded to discrimination testing; therefore, all discrimination testing occurred in Cohort B.

#### Discrimination testing

Testing occurred on Tuesdays and Fridays, with training sessions on Mondays, Wednesdays, Thursdays, as long as the rats met the testing criteria of >80% of responses on the appropriate lever (prior to the first reinforcer and in the total session) in 3 of the last 4 consecutive training sessions. If a rat did not meet the testing criteria, the rat underwent a training session and was not tested. On test days, rats were injected with the dose of the test compound and immediately placed in the operant chamber. Test sessions were identical to training sessions, including a 10-minute timeout before lever insertion, except that both levers were reinforced on an FR10 schedule to prevent bias in lever selection and to allow for response rate determination. Additionally, test sessions were terminated 2 minutes after the first reinforcer was obtained or after 15 minutes if no reinforcer was obtained. Tests for each compound occurred within subjects and the dose order was counterbalanced to prevent the order of testing from influencing results.

### Quantification and statistical analyses

The training data for cohorts A and B were initially analyzed together until they were split into separate groups and training differed. Once separated, all training data for each cohort were again analyzed separately to allow for the visualization and analysis of different training doses of fentanyl. For each training and testing session, the primary measure of discrimination was the percent of fentanyl-appropriate lever presses prior to delivery of the first reinforcer because this represents behavior prior to feedback from sucrose delivery. Full expression of the discriminative effects of fentanyl was defined as >80% fentanyl-appropriate responses prior to the delivery of the first reinforcer. Partial substitution was described as between 40-80% fentanyl-appropriate responses prior to delivery of the first reinforcement (Jaramillo et al. 2016; Solinas et al. 2006). The percentage of total fentanyl-appropriate lever responses, response rate (lever presses / min), and latency to first reinforcer (in seconds) were also calculated. During training and testing sessions, if a rat did not complete an FR10, the percent first could not be calculated and was excluded, but the response rate was calculated and included in the analyses, and reward latency was recorded as 900 seconds (the length of the session). The median effective dose (ED50), the predicted dose that would produce 50% fentanyl-appropriate responding, was calculated using a simple linear regression on the 2 doses which encompass 50% fentanyl-appropriate responding. If no value for a rat was below 50%, a value of 0% was used for a 0 mg/kg dose. If no value for a rat was above 50% for any xylazine co-administration condition (n=3), the rat was excluded. Because there was an apparent bimodal distribution of ED50 in male rats when 0.1 mg/kg xylazine was administered with the dose range of fentanyl, we used F tests to compare variance between males and females, which was performed separately for each xylazine condition (fentanyl + 0, 0.1, or 0.32 mg/kg xylazine). Fisher’s exact tests were used to analyze contingency tables comparing the proportion of males and females that met testing criteria after training at the different fentanyl training doses. All statistical analyses were performed on GraphPad Prism and SPSS Statistics software, and significance was set at p<0.05. Graphs were made using GraphPad Prism software. All data were initially analyzed with sex as an independent variable, and if there was no main effect of sex, data were collapsed across sex. Training data for each fentanyl training dose were analyzed separately, with drug treatment (fentanyl vs saline) as a within-subjects variable and training session as a repeated-measures within-subjects variable. Effects of the 3 training doses of fentanyl in Cohort A on response rate and reward seeking suppression were analyzed using one-way ANOVAs with fentanyl dose as a within-subjects variable. Test data were analyzed with fentanyl dose and/or xylazine dose as within-subjects variables and sex as a between-subjects variable when described. Median effective dose of fentanyl was analyzed using a one-way ANOVA with xylazine dose as a within-subjects variable. When interactions were significant, a Sidak’s or Tukey’s multiple comparisons test was used for post-hoc analysis.

## Results

### Long-Evans rats learn to discriminate 0.01 mg/kg fentanyl from saline, but most do not meet testing criteria

Rats (n=23; 12 females and 11 males) underwent discrimination training followed by testing the discriminative stimulus control of fentanyl, and then xylazine substitution, and effects of xylazine co-administration on fentanyl discrimination (Fig. 1A). Rats were first trained to discriminate 0.01 mg/kg fentanyl from saline. In daily sessions, rats learned that one lever was reinforced with sucrose on an FR10 schedule (fentanyl-appropriate lever) 10 minutes after fentanyl had been administered by the experimenter, and on other days the other lever was reinforced after saline was administered (saline-appropriate lever). Analysis of acquisition of 0.01 mg/kg fentanyl discrimination (sessions 1-42) showed no main effect of sex on the percent fentanyl-appropriate responses (F_(1,757)_=1.506, p=0.2202) (3-way mixed effects analysis), so males and females were combined for additional analyses. The two-way mixed effects analysis showed a main effect of training session (F_(41,902)_=1.797, p=0.0018), fentanyl (F_(1,22)_=287.5, p<0.0001), and an interaction between training session and fentanyl (F_(41,798)_=11.89, p<0.0001; Fig. 1B). Post-hoc analyses indicated that the percent fentanyl-appropriate responses was greater on fentanyl vs. saline sessions in most cases, including sessions 18-42 (ps<0.05). However, in general accuracy performance was below 80% fentanyl-appropriate responding on fentanyl sessions.

**Fig. 1.**
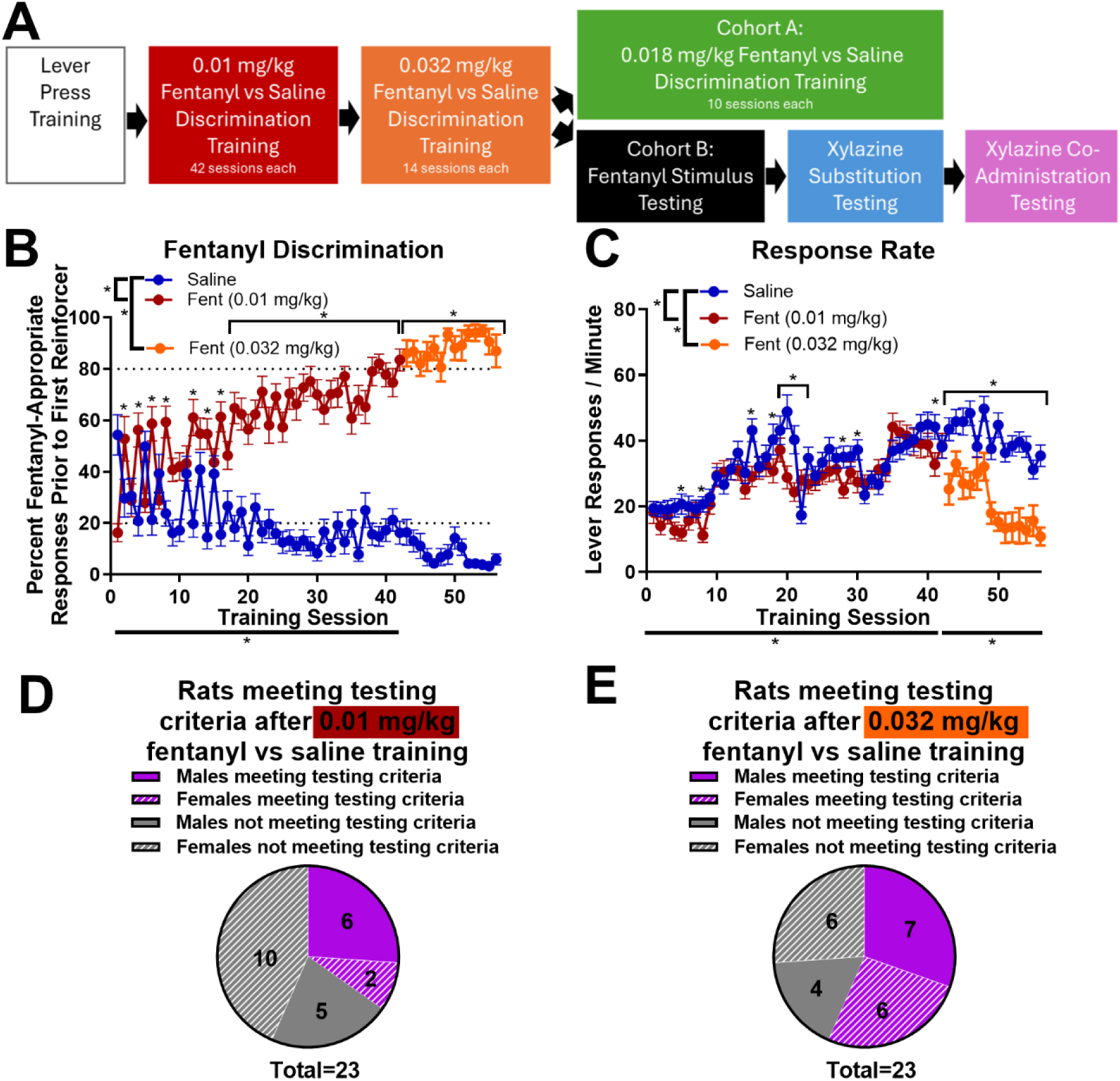
Rats learn to discriminate interoceptive effects of fentanyl from saline. Timeline of experiments (A). All rats (n=23, 12 females and 11 males) were initially trained to discriminate fentanyl from saline in an operant discrimination task. After 42 training sessions each of saline and 0.01 mg/kg fentanyl, the training dose was increased to 0.032 mg/kg. For the percent fentanyl-appropriate responses prior to the first reinforcer, a measure of discrimination accuracy, there was a main effect of 0.01 mg/kg fentanyl compared to saline, a main effect of training session, and a 0.01 mg/kg fentanyl × training session interaction (B). When the training dose was increased to 0.032 mg/kg, there was a main effect of fentanyl and a 0.032 mg/kg fentanyl × training session interaction (B). Post-hoc analyses showed higher percent fentanyl-appropriate responses in several fentanyl sessions, indicated by asterisks. For response rates, there were main effects of fentanyl, training session, and fentanyl × training session interactions for both doses, and post-hoc analyses showed differences in response rates between fentanyl and saline sessions, indicated by asterisks (C). Pie charts express the distribution of rats that met testing criteria after discrimination training at 0.01 mg/kg fentanyl (D) and 0.032 mg/kg fentanyl (E). Black dotted lines at y=20% and 80% indicate boundaries for testing criteria (B). *p<0.05.

Response rate for each session was used as a measure of the effects of fentanyl on locomotor behavior. There was no main effect of sex on response rate (F_(1,21)_=0.5669, p=0.4598) (3-way ANOVA), so males and females were combined for additional analyses. There was a main effect of training session (F_(41,902)_=23.51, p<0.0001), a main effect of 0.01 mg/kg fentanyl (F_(1,22)_=5.808, p=0.0248), and an interaction between training session and fentanyl (F_(41,902)_=7.307, p<0.0001) for response rate (2-way ANOVA; Fig. 1C). Post-hoc analyses indicated that response rate was lower (ps<0.05) on 10 fentanyl sessions and on a single saline session.

Together, these data show that rats were able to discriminate between the effects of 0.01 mg/kg fentanyl and saline. Additionally, 0.01 mg/kg fentanyl produced a minor suppression of response rate on some days. However, after 42 sessions of each session type (fentanyl and saline), few rats (n=8) met the testing criteria that requires consistent percent fentanyl-appropriate responses prior to the first reinforcer above 80% on fentanyl sessions and below 20% on saline sessions, and there was no significant difference between the proportion of males and females that met testing criteria (p=0.0894; Fisher’s exact test; Fig. 1D). Therefore, to enhance discrimination, we increased the training dose of fentanyl to 0.032 mg/kg.

### Higher training doses of fentanyl (0.032 and 0.018 mg/kg) promote discrimination but suppress behavior

Training data for the 0.032 mg/kg fentanyl training vs. saline were analyzed separately (sessions 43-56). For 0.032 mg/kg fentanyl, there was no main effect of sex on the percent fentanyl-appropriate responses (F_(1,205)_=1.410, p=0.2365) (3-way mixed effects analysis), so males and females were combined for additional analyses. There was a main effect of fentanyl (F_(1,22)_=982.0, p<0.0001) and an interaction between training session and fentanyl (F_(13,218)_=2.229, p=0.0094), but no main effect of training session (F_(13,286)_=0.8073, p=0.6523) (2-way mixed effects analysis; Fig. 1B). Post-hoc analyses indicated higher percent fentanyl-appropriate responses on every 0.032 mg/kg fentanyl training session compared to saline training sessions (ps<0.05). Analysis of response rates showed no main effect of sex (F_(1,21)_=0.5058, p=0.4848; 3-way ANOVA), so males and females were combined for additional analyses. There was a main effect of training session (F_(13,286)_=22.11, p<0.0001), a main effect of fentanyl (F_(1,22)_=30.98, p<0.0001), and an interaction between training session and fentanyl (F_(13,286)_=4.869, p<0.0001) for response rate (2-way ANOVA; Fig. 1C). Post-hoc analyses indicated that response rate was lower on every 0.032 mg/kg fentanyl training session compared to saline training sessions (ps<0.05).

After increasing the training dose to 0.032 mg/kg fentanyl for 28 sessions (14 each of fentanyl and saline), most rats (n=13, 6 females and 7 males) met testing criteria, but a subset of rats (n=10, 6 females and 4 males) continued to fail to meet testing criteria due to lack of discrimination or suppression of response rate, though there was no difference between the proportion of males and females that met testing criteria (p=0.6802; Fisher’s exact test; Fig. 1E).

*Cohort A.* In the group of rats that did not meet testing criteria (n=10, 6 females and 4 males; Cohort A), we decreased the training dose to an intermediate dose (0.018 mg/kg) to examine if it would promote discrimination without suppressing response rate. All training data for Cohort A are shown separately in Figure 2 and each training dose was analyzed separately. Males and females were combined for analysis because of the small number of subjects. For 0.01 mg/kg fentanyl (sessions 1-42), there was a main effect of fentanyl (F_(1,9)_=36.23, p=0.0002) and an interaction between training session and fentanyl (F_(41,312)_=5.323, p<0.0001) on the percent fentanyl-appropriate responses, but there was no main effect of training session (F_(41,369)_=1.286, p=0.1195; 2-way mixed effects analysis; Fig. 2A). Post-hoc analyses indicated higher percent fentanyl-appropriate responses on several fentanyl sessions compared to saline sessions (ps<0.05). When the dose was increased to 0.032 mg/kg fentanyl (sessions 43-56), there was a main effect of fentanyl on the percent fentanyl-appropriate responses (F_(1,9)_=383.3, p<0.0001), but no main effect of training session (F_(13,17)_=0.8680, p=0.5882) or interaction between fentanyl and training session (F_(13,62)_=1.043, p=0.4238; 2-way mixed effects analysis; Fig. 2A). Lastly, when the dose was lowered to 0.018 mg/kg fentanyl (sessions 57-66), there was a main effect of fentanyl (F_(1,9)_=141.7, p<0.0001) and an interaction between training session and fentanyl (F_(9,53)_=2.933, p=0.0067) for the percent fentanyl-appropriate responses, but there was no significant main effect of training session (F_(9,81)_=1.804, p=0.0801; 2-way mixed effects analysis; Fig. 2A). Post-hoc analyses indicated higher percent fentanyl-appropriate responses on every 0.018 mg/kg fentanyl training session compared to saline training sessions (ps<0.05).

**Fig. 2.**
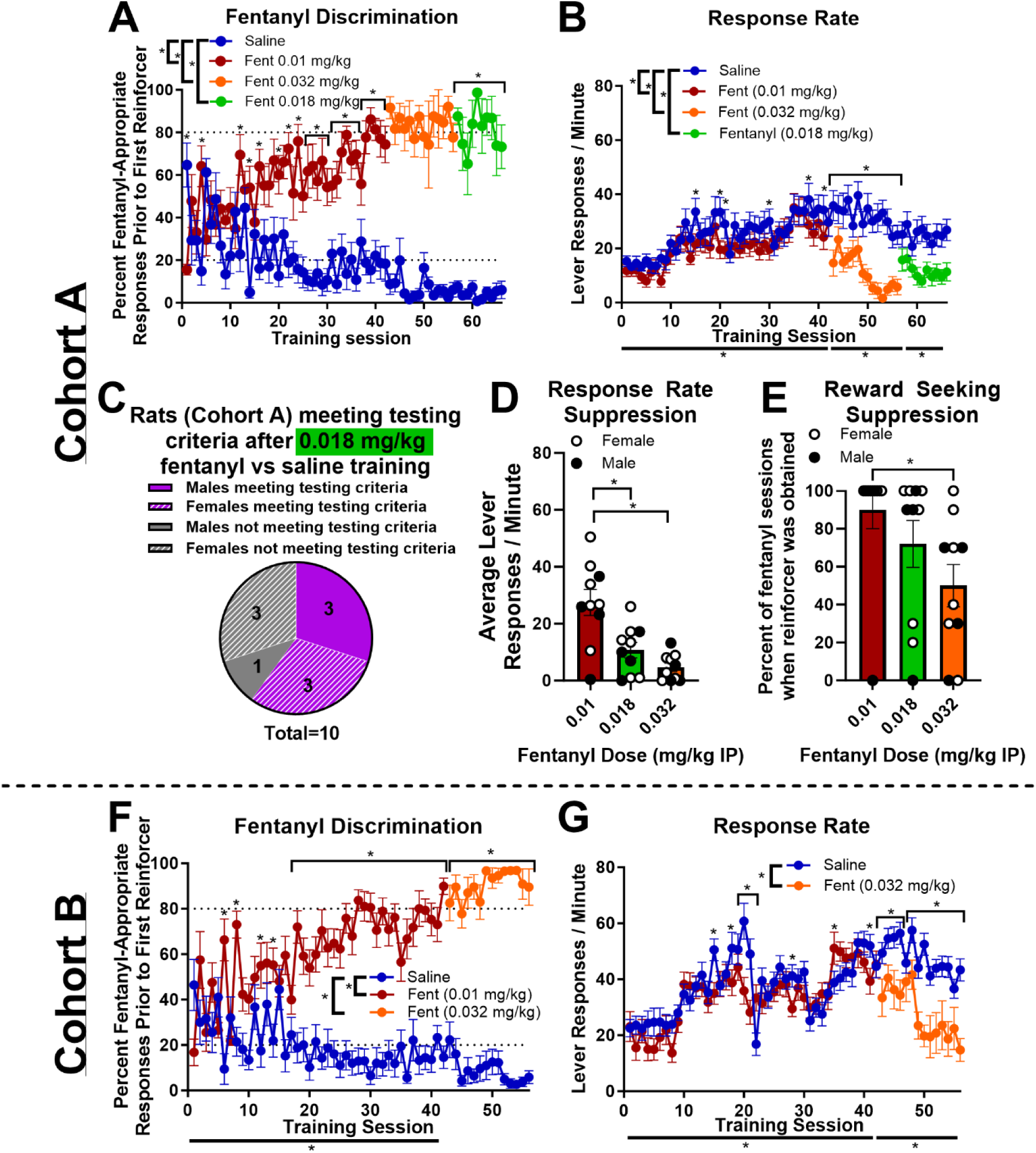
Higher training doses of fentanyl promote discrimination but suppress behavior. Rats that did not meet testing criteria after 0.032 mg/kg fentanyl discrimination training (Cohort A, n=10) were additionally trained to discriminate an intermediate dose of fentanyl (0.018 mg/kg). For the percent fentanyl-appropriate responses prior to the first reinforcer, there was a main effect of fentanyl compared to saline for all three fentanyl doses, and for 0.01 mg/kg and 0.018 mg/kg fentanyl, there was a fentanyl × training session interaction (A). Post-hoc analyses showed higher percent fentanyl-appropriate responses in several fentanyl sessions, indicated by asterisks (A). For both 0.01 and 0.032 mg/kg fentanyl, there were main effects of fentanyl and training session on response rate, and there were also fentanyl × training session interactions, and post-hoc analyses showed differences in response rates between fentanyl and saline sessions, indicated by asterisks (B). For 0.018 mg/kg fentanyl, there was a main effect of fentanyl and a main effect of training session on response rate (B). A pie chart expresses the distribution of rats that met testing criteria after training at 0.018 mg/kg fentanyl (C). When the average response rate of the final 5 fentanyl training sessions was compared for each dose in Cohort A, there was a main effect of fentanyl dose on response rate, and post-hoc analyses showed that response rate was lower for both 0.018 and 0.032 mg/kg fentanyl compared to 0.01 mg/kg (D). When the percent of training sessions in which at least one reinforcer was obtained were compared between the final 10 training sessions at each fentanyl training dose, there was a main effect of fentanyl dose, and post-hoc analyses showed that the percent of sessions when reinforcer was obtained was only significantly reduced for 0.032 mg/kg compared to 0.01 mg/kg fentanyl (E). Open symbols indicate females, and closed symbols indicate males (D-E). Rats that met testing criteria after 0.032 mg/kg fentanyl discrimination training (Cohort B, n=13) proceeded to testing, but their training data were separated and analyzed (F-G). For the percent fentanyl-appropriate responses prior to the first reinforcer, there was a main effect of fentanyl compared to saline and a fentanyl × training session interaction for both training doses, but there was only a main effect of training session for 0.01 mg/kg fentanyl (F). Post-hoc analyses showed higher percent fentanyl-appropriate responses in several fentanyl sessions, indicated by asterisks (F). For both 0.01 and 0.032 mg/kg fentanyl, there were main effects of fentanyl and training session on response rate, and there were also fentanyl × training session interactions, and post-hoc analyses showed differences in response rates between fentanyl and saline sessions, indicated by asterisks (G). Black dotted lines at y=20% and 80% indicate boundaries for testing criteria (B, F). *p<0.05.

Response rates were analyzed similarly to fentanyl-appropriate responses. At the 0.01 mg/kg fentanyl dose (sessions 1-42), there was a main effect of fentanyl (F_(1,9)_=5.420, p=0.0449), a main effect of training session (F_(41,369)_=10.10, p<0.0001), and a fentanyl by training session interaction (F_(41,369)_=2.155, p<0.0001; 2-way ANOVA; Fig. 2B). Post-hoc analyses indicated that response rate was lower on 6 0.01 mg/kg fentanyl sessions compared to saline sessions (ps<0.05). When the dose was increased to 0.032 mg/kg fentanyl (sessions 43-56), there was a main effect of fentanyl (F_(1,9)_=33.78, p=0.0003), a main effect of training session (F_(13,117)_=7.208, p<0.0001), and an interaction between fentanyl and training session (F_(13,117)_=3.201, p=0.0004; 2-way ANOVA; Fig. 2B). Post-hoc analyses indicated that response rate was lower in all 0.032 mg/kg fentanyl sessions compared to saline sessions (ps<0.05). When the dose was lowered to 0.018 mg/kg (sessions 57-66), there was a main effect of fentanyl (F_(1,9)_=17.03, p=0.0026) and a main effect of training session (F_(9,81)_=2.724, p=0.0079), but no interaction between fentanyl and training session (F_(9,81)_=1.860, p=0.0699; 2-way ANOVA; Fig. 2B).

In Cohort A, more than half of the rats that failed to meet testing criteria after 0.032 mg/kg fentanyl training later met testing criteria when the dose was lowered to 0.018 mg/kg fentanyl training, and there was no difference between the proportion of males and females that met testing criteria (p=0.5714; Fisher’s exact test; Fig. 2C). Together, these data suggest that higher training doses of fentanyl produced more effective discrimination in rats that continued performing the task, but also suppressed response rates. To compare response rate suppression across the three training doses of fentanyl, the average response rate of the final 5 fentanyl training days was compared for each dose, and there was a main effect of fentanyl dose on response rate (F_(2,18)_=36.90, p<0.0001) (one-way repeated measures ANOVA) (Fig. 2D). Post-hoc analyses indicated that response rate was lower for both 0.018 mg/kg fentanyl (p<0.0001) and 0.032 mg/kg fentanyl (p<0.0001) compared to 0.01 mg/kg fentanyl, but there was no difference between 0.018 mg/kg and 0.032 mg/kg (p=0.0994). In some cases, response rates were so greatly suppressed that rats failed to obtain a single reinforcer across several fentanyl sessions. To quantify this, we calculated the percent of training sessions in which at least one reinforcer was obtained for the final 10 fentanyl sessions at each training dose, and there was a main effect of fentanyl dose (F_(2,18)_=7.933, p=0.0034; one-way RM ANOVA; Fig. 2E). Post-hoc analyses indicated that the percentage of sessions when reinforcer was obtained was reduced for 0.032 mg/kg fentanyl compared to 0.01 mg/kg fentanyl (p=0.0024), but there was no difference between 0.01 and 0.018 (p=0.2013) or between 0.018 and 0.032 (p=0.1007). These results suggest that although 0.018 mg/kg fentanyl does suppress response rate, it doesn’t significantly prevent subjects’ ability to complete the task and may therefore be a more appropriate training dose. Due to their complex training history, Cohort A did not proceed to testing.

*Cohort B.* The rats that met testing criteria after 0.032 mg/kg fentanyl vs. saline discrimination training (n=13, 6 females and 7 males; Cohort B) were used for the remainder of the experiments, and their training data were separated and each training dose was analyzed separately. There was no main effect of sex on percent fentanyl-appropriate responses for either 0.01 mg/kg (sessions 1-42) fentanyl (F_(1,402)_=1.811, p=0.1792) or 0.032 mg/kg (sessions 43-56) fentanyl (F_(1,40)_=0.1466, p=0.7038; 3-way mixed effects analyses), and there was also no main effect of sex on response rate for either 0.01 mg/kg fentanyl (F_(1,11)_=4.595, p=0.0553) or 0.032 mg/kg fentanyl (F_(1,11)_=0.1168, p=0.1168; 3-way ANOVAs). Therefore, males and females were combined for analyses. At the 0.01 mg/kg training dose (sessions 1-42), there was a main effect of fentanyl (F_(1,12)_=331.6, p<0.0001), a main effect of training session (F_(41,496)_=1.601, p=0.0122), and an interaction between fentanyl and training session (F_(41,443)_=7.352 p<0.0001) for the percent fentanyl-appropriate responses (2-way mixed-effects analysis; Fig. 2F). Post-hoc analyses indicated that the percent fentanyl-appropriate responses was greater in fentanyl sessions than saline sessions in most cases, including sessions 18-42 (ps<0.05). When the training dose was increased to 0.032 mg/kg (sessions 43-56), there was a main effect of fentanyl (F_(1,12)_=515.5, p<0.0001) and an interaction between fentanyl and training session (F_(13,143)_=2.537. p=0.0036), but no main effect of training session (F_(13,156)_=1.083, p=0.3780; 2-way mixed-effects analysis; Fig. 2F). Post-hoc analyses indicated higher percent fentanyl-appropriate responses on every 0.032 mg/kg fentanyl training session compared to saline training sessions (ps<0.05).

For response rate, there was a main effect of training session (F_(41,492)_=15.63, p<0.0001) and an interaction between 0.01 mg/kg fentanyl and training session (F_(41,492)_=6.111, p<0.0001), but no main effect of 0.01 mg/kg fentanyl (F_(1,12)_=1.819, p=0.2024; 2-way ANOVA; Fig. 2G). Post-hoc analyses indicated that response rate was lower in 6 0.01 mg/kg fentanyl sessions and higher in 2 fentanyl sessions compared to saline sessions (ps<0.05). There was a main effect of 0.032 mg/kg fentanyl (F_(1,12)_=11.16, p=0.0059), a main effect of training session, (F_(13,156)_=15.38, p<0.0001), and an interaction between fentanyl and training session (F_(13,156)_=2.964, p=0.0007) for response rate (2-way ANOVA; Fig. 2G). Post-hoc analyses indicated that response rates were lower on all but one 0.032 mg/kg fentanyl training session compared to saline session (ps<0.05). Together, these data suggest that 0.032 mg/kg fentanyl suppressed response rates, but rats were able to discriminate effectively between this dose of fentanyl and saline. Additionally, as a group, when the dose was increased to 0.032 mg/kg, accuracy performance was at or above 80% on fentanyl sessions and below 20% on saline sessions.

### Lever choice is controlled by the discriminative stimulus effects of fentanyl (Cohort B)

After rats met testing criteria for 0.032 mg/kg fentanyl discrimination (Cohort B, n=13; 7 males and 6 females), they underwent a series of test sessions interspersed with training sessions. Four doses of fentanyl were tested to confirm discriminative stimulus control of fentanyl. There was no main effect of sex on the percent fentanyl-appropriate responses (F_(1,11)_=0.5132, p=0.4887), response rate (F_(1,11)_=0.6616, p=0.4333), or reinforcer latency (F_(1,11)_=0.2296, p=0.6412) (2-way mixed-effects analysis and 2-way ANOVAs), so males and females were combined for additional analyses but are shown separately on graphs for illustrative purposes.

Percent fentanyl-appropriate responses increased with increasing doses of fentanyl, which was demonstrated by a main effect of fentanyl dose (F_(3,33)_=98.12, p<0.0001) on the percent fentanyl-appropriate responses (one-way mixed effects analysis) (Fig. 3A). Post-hoc analyses indicated that percent fentanyl-appropriate responses significantly increased for all fentanyl doses compared to 0.0032 mg/kg (0.005 mg/kg p=0.0156; 0.018 and 0.032 mg/kg p<0.0001). Additionally, the average percent fentanyl-appropriate responses for the 0.018 and 0.032 mg/kg fentanyl doses was greater than 80% for both males and females. Therefore, these results suggest that 0.018 mg/kg fully substituted for the 0.032 mg/kg training dose. The lower fentanyl doses (0.0032 mg/kg and 0.005 mg/kg) averaged below 40%, and therefore did not substitute for the training dose. These data suggest that lever choice was guided by the interoceptive effects of fentanyl.

**Fig. 3.**
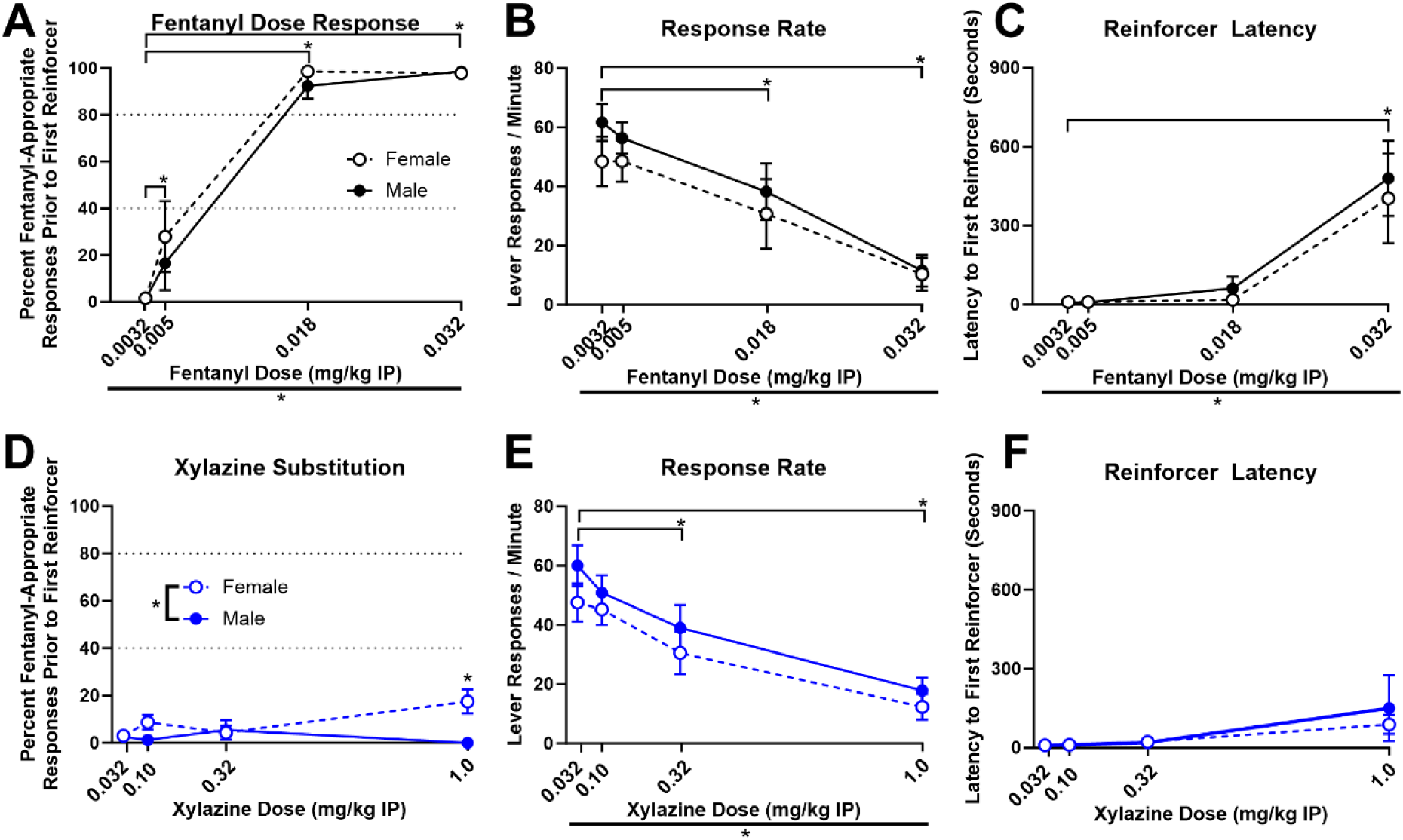
Xylazine does not substitute for the interoceptive effects of fentanyl despite promoting similar reductions in response rate. Rats from cohort B (n=13, 6 females and 7 males) proceeded to discrimination testing. A dose range of fentanyl was used to test discriminative stimulus control of fentanyl. Females (open symbols) and males (closed symbols) were combined for analysis unless there was a main effect of sex. There was a main effect of fentanyl dose on percent fentanyl-appropriate responses (A), response rate (B), and reinforcer latency (C), a measure of the time elapsed between lever insertion (session start) and the first FR10 completion. Post-hoc analyses showed that compared to the lowest 0.0032 mg/kg dose, percent fentanyl-appropriate responses were greater for all higher doses (A), response rate was lower for 0.018 and 0.032 mg/kg fentanyl doses (B), and reinforcer latency was higher for 0.032 mg/kg fentanyl (C). A dose range of xylazine was administered alone in place of fentanyl to evaluate its ability to substitute for fentanyl. There was a main effect of sex on percent fentanyl-appropriate responses after xylazine administration, and post-hoc analyses showed that females made a higher percent fentanyl-appropriate responses compared to males after administration of 1 mg/kg xylazine (D). There was a main effect of xylazine dose on response rate, and post-hoc analyses showed that compared to the lowest 0.032 mg/kg xylazine dose, response rate was reduced for 0.32 and 1 mg/kg xylazine (E). There was no main effect of xylazine dose on reinforcer latency (F). Black dotted line at y=80% indicates minimum cutoff for full substitution, and gray dotted line at y=40% indicates minimum cutoff for partial substitution (A, D). *p<0.05.

Response rate decreased as fentanyl dose increased, which was indicated by a main effect of fentanyl dose on response rate (F_(3,36)_=40.65, p<0.0001; one-way RM ANOVA; Fig. 3B). Post-hoc analyses indicated that compared to 0.0032 mg/kg fentanyl, response rate was significantly reduced for 0.018 (p=0.0002) and 0.032 mg/kg (p<0.0001), but not for 0.005 mg/kg (p=0.8733). Because we noticed that higher doses of fentanyl suppressed the initiation of reward seeking, we also measured reinforcer latency, the time elapsed between lever insertion (session start) and the first reinforcer delivery. There was a main effect of fentanyl dose (F_(3,36)_=16.15, p<0.0001) on reinforcer latency (one-way RM ANOVA; Fig. 3C), and post-hoc analyses indicated that compared to 0.0032 mg/kg fentanyl, reinforcer latency was significantly longer for 0.032 mg/kg (p<0.0001) but was unchanged for 0.005 (p>0.999) and 0.018 (p=0.9456). Together, these effects of fentanyl on response rate and reinforcer latency indicate that increasing doses of fentanyl increasingly suppressed behavior.

### Xylazine does not substitute for fentanyl despite promoting similar reductions in response rate (Cohort B)

Xylazine was administered in place of fentanyl to determine if xylazine alone would substitute for the interoceptive effects of fentanyl. When xylazine alone was administered, there was a main effect of sex (F_(1,11)_=7.707, p=0.0180) and an interaction between xylazine dose and sex (F_(3,32)_=4.347, p=0.0112) on the percent fentanyl-appropriate responses prior to the first reinforcer, but there was no main effect of xylazine dose (F_(3,32)_=1.463, p=0.2430; 2-way mixed effects analysis; Fig. 3D). Post-hoc analyses indicated that females made a higher percentage of fentanyl-appropriate responses than males after administration of 1 mg/kg xylazine (p=0.0006). The average percent fentanyl-appropriate responses for all doses of xylazine and for both sexes was below 40%, which indicates that the interoceptive effects of xylazine at any dose tested did not substitute for fentanyl.

There was no main effect of sex on response rate (F_(1,11)_=1.289, p=0.2804) or reinforcer latency (F_(1,11)_=0.1796, p=0.6799; 2-way ANOVAs), so males and females were combined for additional analyses but are shown separately on graphs for illustrative purposes. Response rate decreased as xylazine dose increased, which was indicated by a main effect of xylazine dose on response rate (F_(3,36)_=35.47, p<0.0001; one-way RM ANOVA; Fig. 3E). Post-hoc analyses indicated that compared to 0.032 mg/kg xylazine, response rate was significantly reduced for 0.32 (p=0.0001) and 1 mg/kg (p<0.0001) but was not different for 0.1 mg/kg (p=0.3412). There was no main effect of xylazine dose on reinforcer latency (F_(3,36)_=2.549, p=0.0710; one-way RM ANOVA; Fig. 3F). Together, these data suggest that xylazine does not produce similar interoceptive effects of fentanyl despite having similar sedative effects on response rate.

### Xylazine potentiates fentanyl discrimination and promotes partial substitution of lower fentanyl doses (Cohort B)

Two xylazine doses (0.1 and 0.32 mg/kg) were combined with the four fentanyl doses and compared to fentanyl alone to determine if xylazine altered interoceptive sensitivity to fentanyl. There was no main effect of sex on the percent fentanyl-appropriate responses (F_(1,11.774)_=0.131, p=0.724; 3-way mixed effects analysis), so animals were collapsed across sex for concise visualization. There was a main effect of fentanyl dose (F_(3,36)_=51.74, p<0.0001) and an interaction between fentanyl dose and xylazine dose (F_(6,66)_=4.504, p=0.0007) on the percent fentanyl-appropriate responses, but there was no main effect of xylazine dose (F_(2,24)_=2.623, p=0.0933; 2-way mixed effects analysis; Fig. 4A). Post-hoc analyses indicated that for the lowest fentanyl dose (0.0032 mg/kg), the addition of 0.32 mg/kg xylazine significantly increased the percent fentanyl-appropriate responses compared to fentanyl alone (p=0.0020) or 0.1 mg/kg xylazine + fentanyl (p=0.0055). Additionally, the average percent fentanyl-appropriate responses was 40% or greater when 0.32 mg/kg xylazine was added to either of the lowest fentanyl doses (0.0032 and 0.005 mg/kg), which suggests partial substitution for the fentanyl training dose and indicates potentiation of the interoceptive effects of the lower fentanyl doses. Interestingly, the average percent fentanyl-appropriate responses was also below 80% when either 0.1 or 0.32 mg/kg xylazine were added to the highest fentanyl doses (0.018 and 0.032 mg/kg), which suggests partial substitution, an indication that xylazine may blunt or alter the interoceptive effects of higher doses of fentanyl.

**Fig. 4.**
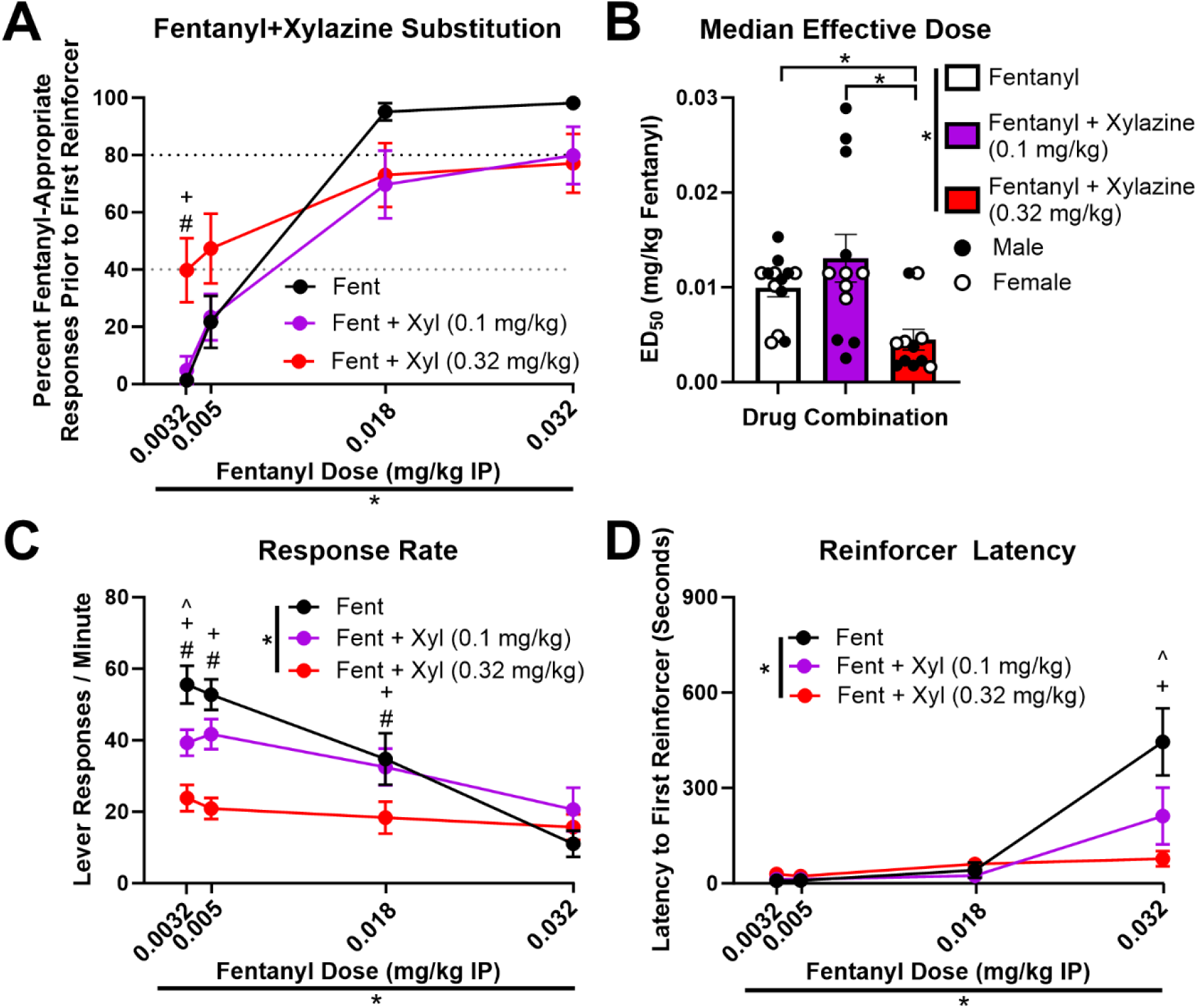
Xylazine potentiates interoceptive effects of fentanyl and promotes partial substitution of lower fentanyl doses. In a series of tests, 2 doses of xylazine were combined with the dose range of fentanyl to examine the effects of xylazine on fentanyl interoception. There was a main effect of fentanyl dose and a fentanyl dose × xylazine dose interaction for the percent fentanyl-appropriate responses, and post-hoc analyses showed that the addition of 0.32 mg/kg xylazine to 0.0032 mg/kg fentanyl significantly increased the percent fentanyl-appropriate responses compared to fentanyl alone or fentanyl + 0.1 mg/kg xylazine (A). There was a main effect of xylazine dose on ED50, and post-hoc analyses showed that ED50 was significantly reduced when 0.32 mg/kg xylazine was co-administered with fentanyl compared to fentanyl alone or fentanyl + 0.1 mg/kg xylazine (B). There were main effects of fentanyl dose and xylazine dose as well as fentanyl dose × xylazine dose interactions for both response rate (C) and reinforcer latency (D). For both response rate and reinforcer latency, post-hoc analyses showed differences between xylazine groups at some fentanyl doses, which are indicated by symbols. ^p<0.05 for fentanyl alone vs. fentanyl + 0.1 mg/kg xylazine. +p<0.05 for fentanyl alone vs. fentanyl + 0.32 mg/kg xylazine. #p<0.05 for fentanyl + 0.1 mg/kg xylazine vs. fentanyl + 0.32 mg/kg xylazine. Black dotted line at y=80% indicates minimum cutoff for full substitution, and gray dotted line at y=40% indicates minimum cutoff for partial substitution (A). *p<0.05.

Percent fentanyl-appropriate responses were used to calculate the ED50 for each rat for fentanyl alone, fentanyl + 0.1 mg/kg xylazine, and fentanyl + 0.32 mg/kg xylazine. There was no main effect of sex on the ED50 of fentanyl (F_(1,11)_=0.3658, p=0.5576; 2-way mixed effects analysis), so animals were collapsed across sex. There was a main effect of xylazine dose on ED50 (F_(2,21)_=9.621, p=0.0011; one-way mixed effects analysis; Fig. 4B), and post-hoc analyses indicated that the median effective dose was lower for fentanyl + 0.32 mg/kg xylazine compared to fentanyl alone (p=0.0343) or fentanyl + 0.1 mg/kg xylazine (p=0.0008). Together, these results suggest that the addition of 0.32 mg/kg xylazine potentiated the interoceptive effects of the lowest fentanyl doses (0.0032 and 0.005 mg/kg) and the addition of either 0.1 or 0.32 mg/kg xylazine slightly blunted the interoceptive effects of the higher fentanyl doses (0.018 and 0.032 mg/kg). Additionally, although there was no main effect of sex on ED50, we did notice an apparent bimodal distribution in males when 0.1 mg/kg xylazine was co-administered with fentanyl, where some rats showed an increase. To compare the variability between males and females separately at each dose of xylazine co-administration, we performed F tests. When 0.1 mg/kg xylazine was co-administered with fentanyl, there was a significant difference in the variance between males and females (F_(6,4)_=92.06, p=0.0006; F test to compare variance). There was no difference in the variance between males and females for the ED50 of fentanyl with no xylazine (F_(5,6)_=1.037, p=0.9469) or with 0.32 mg/kg xylazine (F_(5,4)_=1.052, p=0.9881; F test). Therefore, the effects of 0.1 mg/kg xylazine on fentanyl interoception are more variable in males.

### Xylazine potentiates fentanyl-induced suppression of response rates and locomotor rates but reduces suppression of initiation of reward seeking (Cohort B)

There was no main effect of sex on response rate (F_(1,11)_=1.381, p=0.265, 3-way ANOVA), so animals were collapsed across sex. There were main effects of fentanyl dose (F_(3,36)_=31.56 p<0.0001) and xylazine dose (F_(2,24)_=29.78, p<0.0001) on response rate, and there was also an interaction between fentanyl dose and xylazine dose (F_(6,72)_=7.746, p<0.0001; 2-way ANOVA; Fig. 4C). Post-hoc analyses indicated that for 0.0032 mg/kg fentanyl, the response rate was lower than fentanyl alone when fentanyl was combined with 0.1 (p=0.0065) or 0.32 mg/kg xylazine (p<0.0001), and response rate was lower with the addition of 0.32 compared to 0.1 mg/kg xylazine (p=0.0119). Additionally, for 0.005 mg/kg fentanyl, the response rate was lower than fentanyl alone when fentanyl was combined with 0.32 mg/kg xylazine (p<0.0001), and response rate was lower with the addition of 0.32 compared to 0.1 mg/kg xylazine (p=0.0002). Finally, for 0.018 mg/kg fentanyl, response rate was lower than fentanyl alone when fentanyl was combined with 0.32 mg/kg xylazine (p=0.0059), and response rate was lower with the addition of 0.32 compared to 0.1 mg/kg xylazine (p=0.0287). These results suggest that the addition of xylazine to fentanyl exacerbated the suppressive effects of fentanyl on response rate.

There was no main effect of sex on reinforcer latency (F_(1,11)_=0.175, p=0.684, 3-way ANOVA), so animals were collapsed across sex. There were main effects of fentanyl dose (F_(3,36)_=21.40, p<0.0001) and xylazine dose (F_(2,24)_=4.290, p=0.0255) on reinforcer latency, and there was also an interaction between fentanyl dose and xylazine dose (F_(6,72)_=5.737, p<0.0001; 2-way ANOVA; Fig. 4D). Post-hoc analyses indicated that the addition of 0.1 (p=0.004) or 0.32 mg/kg xylazine (p<0.0001) to 0.032 mg/kg fentanyl significantly reduced reinforcer latency. Therefore, xylazine reduced the suppressive effects of 0.032 mg/kg fentanyl on the initiation of reward seeking.

## Discussion

In the present study, we evaluated the effects of xylazine co-administration on sensitivity to the interoceptive effects of fentanyl in male and female rats trained to discriminate between fentanyl (dose) and vehicle. First, we showed how various fentanyl training doses impacted fentanyl discrimination training and response rate. Then, in a subset of rats that met stringent testing criteria, we confirmed that lever choice was controlled by the interoceptive effects of fentanyl by testing percent responding on the fentanyl-appropriate lever after a range of fentanyl doses. Xylazine administered alone did not increase percent responding on the fentanyl-appropriate lever, which suggests it did not produce fentanyl-like interoceptive effects; however, xylazine had response rate-suppression effects similar to those of fentanyl. Interestingly, the addition of 0.32 mg/kg xylazine to fentanyl significantly increased fentanyl-appropriate responding at the lowest dose of fentanyl and lowered the median effective dose of fentanyl. Overall, these results suggest that xylazine co-administration with fentanyl potentiates sensitivity to the interoceptive effects of fentanyl, particularly by enhancing the interoceptive effects of low doses of fentanyl.

### Acquisition of fentanyl discrimination: manipulation of training doses

A major strength of this work is the inclusion of training data at multiple fentanyl training doses, which can inform future experiments. We began by training rats to discriminate 0.01 mg/kg fentanyl from saline (I.P., 10-minute pretreatment). However, only about a third of rats met testing criteria after over 80 total training sessions. Other work found acquisition of the discrimination using the same fentanyl training dose, pretreatment time, and route of administration in male Sprague Dawley rats, though testing criteria were different (Flynn and France 2021; 2022). As such, some possible explanations for the discrepancy could be the rat strain (Long Evans vs. Sprague Dawley), the inclusion of females, and the more stringent testing criteria in the present study. Given that, on average, rats were responding significantly more on the fentanyl-appropriate lever prior to the first reinforcer on fentanyl days compared to saline days, we theorized that rats were learning to discriminate but were unable to distinguish between this dose of fentanyl and saline sufficiently to meet testing criteria. There is precedence in the drug discrimination field to alter the training dose after initial training (Craft and Stratmann 1996; Green and Grant 1998; Mariathasan et al. 1996; Overton 1979; Stolerman et al. 2011). Therefore, instead of changing our *a priori* defined testing criteria, we decided to increase the training dose to 0.032 mg/kg fentanyl, a dose that was used in another drug discrimination study administered I.P. with a 15-minute pretreatment time (AlKhelb et al. 2022).

Increasing the training dose to 0.032 mg/kg fentanyl increased accuracy and the number of rats that met testing criteria but suppressed response rates and reward seeking. This balance between improved discrimination and suppressed behavior when increasing the training dose has been previously described (Stolerman et al. 2011). In rats that still did not meet testing criteria after training at 0.032 mg/kg (cohort A), we tried a third, intermediate training dose (0.018 mg/kg) to inform future studies. Although response rates were still suppressed compared to 0.01 mg/kg, the percentage of fentanyl sessions when a reinforcer was obtained was not significantly affected. Therefore, we propose that in future studies, it may be advantageous to begin training at 0.01 mg/kg fentanyl to maximize responding during early training, and then to increase the dose to the desired final training dose.

### Discriminative stimulus control of fentanyl

In rats that met testing criteria after 0.032 mg/kg fentanyl training (cohort B), we confirmed discriminative stimulus control of fentanyl on lever choice behavior by examining fentanyl-appropriate responding after administration of a dose range of fentanyl in separate tests. Percent responses on the fentanyl-appropriate lever increased as dose increased, and both the training dose (0.032 mg/kg) and a lower dose this cohort had no prior experience with (0.018 mg/kg) produced robust (>80%) fentanyl-appropriate responding. Additionally, response rate decreased as fentanyl dose increased. Both of these findings are consistent with the fentanyl discrimination literature (AlKhelb et al. 2022; Colpaert et al. 1980; Flynn and France 2021; 2022; Schwienteck et al. 2019).

Fentanyl is an agonist at mu, delta, and kappa opioid receptors, and previous studies have shown that pretreatment with the opioid antagonists naloxone or naltrexone, which have high affinity for the mu opioid receptor, inhibits fentanyl discrimination (AlKhelb et al. 2022; Colpaert et al. 1976b; 1980; Flynn and France 2021; Gharagozlou et al. 2006; Ricarte et al. 2021; Volpe et al. 2011; Yeadon and Kitchen 1988). Notably, naloxone and naltrexone do have effects at other opioid receptors, including kappa opioid receptors (Bianchi and Panerai 1993). Administration of relatively higher doses of other less potent opioid agonists, including heroin and morphine, produce fentanyl-like effects, and low doses of fentanyl produce effects similar to other opioid agonists (AlKhelb et al. 2022; Colpaert et al. 1976a; Colpaert et al. 1978; Craft et al. 1996; Flynn and France 2021; 2022; Schwienteck et al. 2019). In previous studies, both MR 2034, a kappa and mu receptor agonist and the selective delta opioid agonist D-Ala^2^,D-Leu^5^-enkephalin (DADL) substituted for fentanyl interoceptive effects (Emmett-Oglesby and Herz 1987; Shearman and Herz 1982). Together, these data suggest that the interoceptive effects of fentanyl rely on opioid receptor agonism, and the mu and delta receptors both likely play a role, although the role of kappa receptors is less clear.

### Xylazine does not produce fentanyl-like effects

Once discriminative stimulus control of fentanyl was confirmed, we sought to determine if xylazine produced fentanyl-like effects. Xylazine was classically described as an α-2 adrenergic receptor agonist (Kitano et al. 2019; Schwartz and Clark 1998), but our recent evidence indicates xylazine is also a functional agonist of kappa opioid receptors and sigma receptors (Bedard et al. 2024), which interact with opioid receptors (Pasternak 2017). In rats trained to discriminate xylazine, the α-2 adrenergic receptor agonist clonidine fully substituted for the interoceptive effects of xylazine, whereas the α-2 adrenergic receptor antagonist idazoxan blocked xylazine discrimination, which suggests an important role of the α-2 adrenergic receptor agonist effects in the interoceptive effects of xylazine (Colpaert and Janssen 1985). Based on previous literature showing that neither the α-2 adrenergic receptor agonist clonidine nor the selective kappa agonists U69,593 or U50,488 produced morphine-like effects (Craft et al. 1996; Hughes et al. 1996; Picker et al. 1990), we did not expect xylazine alone to have fentanyl-like effects in this study. Indeed, despite causing a suppression of response rate similar to fentanyl, xylazine administration did not significantly increase the percentage of fentanyl-appropriate responding even at the highest dose tested (1 mg/kg). Although it is possible that higher doses of xylazine could produce fentanyl-like effects, when we attempted to test a higher dose of xylazine (3.2 mg/kg), locomotion was severely impaired, and rats were unable to perform the task (data not shown). Interestingly, there is evidence that kappa agonists produce some morphine-like effects in rats trained to discriminate a low dose of morphine (3 mg/kg) but not in rats trained to discriminate 10 mg/kg morphine (Picker et al. 1990). In one study, 10 but not 3.2 mg/kg morphine substituted for the interoceptive effects of 0.032 mg/kg fentanyl, which suggests that the training dose of fentanyl we used in this study may correspond to a higher morphine training dose (AlKhelb et al. 2022). Therefore, it is possible that xylazine could produce fentanyl-like effects in rats trained to discriminate a lower dose of fentanyl. It may also be important to consider our previous finding that withdrawal from morphine dependence can alter the function of α-2 adrenergic receptors in Sprague-Dawley rats, so the effects of xylazine could be different in a model of dependence (McElligott et al. 2013).

### Xylazine effects on fentanyl discrimination

We next sought to determine if xylazine co-administration with fentanyl impacts fentanyl discrimination. When xylazine was administered with fentanyl, an increase in fentanyl-appropriate responding was observed. Specifically, 0.32 mg/kg xylazine+0.0032 mg/kg fentanyl resulted in a significant increase in fentanyl-appropriate responding and responding was at or greater than 40% for both 0.0032 and 0.005 mg/kg fentanyl. Additionally, the median effective dose of fentanyl was significantly reduced. Together, these data suggest that xylazine enhanced discrimination of lower doses of fentanyl that did not produce fentanyl-appropriate responding alone. Rats that undergo this type of discrimination training usually exhibit quantal responding, a phenomenon by which they usually make close to 0% or 100% of their responses on the drug-appropriate lever prior to the first reinforcer (Solinas et al. 2006). Indeed, the increased average percent responding on the fentanyl lever when xylazine was co-administered with fentanyl was primarily driven by full substitution (>80% fentanyl-appropriate responding) in some rats, whereas others responding primarily on the saline-appropriate lever (i.e., <20% fentanyl-appropriate responding). Further studies would be required to determine if the xylazine-induced enhancement of fentanyl discrimination is a stable trait in subjects and the biological factors underlying these individual differences.

Additionally, although the addition of xylazine to 0.018 and 0.032 mg/kg fentanyl did not significantly reduce the percentage of fentanyl-appropriate responding, it did reduce the average to below 80%, which suggests the interoceptive effects of this drug combination do not fully substitute for the interoceptive effects of the training dose of fentanyl (Solinas et al. 2006). This nonsignificant reduction could be caused by the drug combination producing interoceptive effects subjectively different from high doses of fentanyl alone, or it could be due to a leftward shift in the median effective dose, whereby the combined effects of xylazine and fentanyl are much stronger than the training dose of fentanyl.

The mechanism by which xylazine enhances fentanyl discrimination remains to be determined. Interestingly, there are several clinical reports of misuse of clonidine, another α-2 adrenergic receptor agonist, in conjunction with other drugs, including the opioids methadone and heroin as well as benzodiazepines (Beuger et al. 1998; Conway and Balson 1993; Dennison 2001; Lauzon 1992; Schaut and Schnoll 1983; Schindler et al. 2013). Some of these studies indicated that people used clonidine to enhance or prolong the effects of opioids (Beuger et al. 1998; Dennison 2001; Lauzon 1992). Therefore, it is possible that the effects of xylazine at α-2 adrenergic receptors could be contributing to its potentiation of the interoceptive effects of fentanyl. However, preclinical studies have shown that neither the α-2 adrenergic receptor agonists lofexidine or clonidine or antagonists atipamezole or yohimbine impacted morphine discrimination in rats (Hughes et al. 1996; Obeng et al. 2024). Therefore, the role of α-2 adrenergic receptors is unclear and remains to be determined. Because xylazine also has apparent agonist activity at kappa opioid receptors and sigma receptors, it is possible that the effects of xylazine on kappa and sigma receptors could contribute to its enhancement of fentanyl discrimination (Bedard et al. 2024). Additionally, one study showed that α-2 adrenergic receptor agonists, including xylazine, may interact with the opioid system by displacing opioid radioligand binding at opioid receptors (Browning et al. 1982). Therefore, it is possible that xylazine could indirectly enhance the effects of fentanyl on mu and delta opioid receptors by displacing fentanyl bound to kappa receptors, but further experimentation would be required to confirm this theory. Additionally, the mechanism could be even less direct, where the effects of fentanyl and xylazine on different neural circuits, based on their unique pharmacology, could be interacting, which could be evaluated by circuit-based interventions.

### Effects of fentanyl and xylazine on reinforcer latency

One novel finding from these experiments was that the highest dose of fentanyl (the 0.032 mg/kg training dose) significantly increased the latency to the first reinforcer, or the time it took rats to complete the first FR10 schedule after lever insertion. This was unexpected, given that this increase in latency to reinforcer was not described in another study that used the same dose and route of administration (AlKhelb et al. 2022). However, this previous study employed a cumulative dose test design and a 15-min pretreatment time compared to our 10-min pretreatment, so it is possible suppression of reward seeking was less pronounced after an additional 5 minutes of pretreatment or as a result of cumulative testing (AlKhelb et al. 2022). One other study showed that 0.04 mg/kg fentanyl (subcutaneous, 30-min pretreatment) slightly increased latency to reinforcer by an average of a few seconds (Colpaert et al. 1980). In the present study, I.P. administration and a shorter pretreatment time could have exacerbated this effect.

Increased latency to the first reinforcer could be due to either response rate suppression, reduced motivation for sucrose, or a combination of these factors. The fentanyl dose that significantly increased reinforcer latency (0.032 mg/kg) was greater than doses (0.004-0.02 mg/kg) that produced conditioned place preference (CPP) when administered subcutaneously in rats (Gaulden et al. 2021; Solinas et al. 2006), which suggests that 0.032 mg/kg fentanyl injections are reinforcing in rats. Therefore, rats could be less motivated to work for sucrose after administration of a highly reinforcing dose of fentanyl. However, this would contradict previous findings that opioids either do not impact or may increase feeding and sucrose seeking (Castro et al. 2021; Cone et al. 2023; Morley et al. 1984; Nogueiras et al. 2012; Tapia et al. 2019). Interestingly, the addition of 0.32 mg/kg xylazine to 0.032 mg/kg fentanyl abolished the effect of fentanyl on reinforcer latency despite similarly suppressing response rate, so it is unlikely that suppressed responding drove the effect of fentanyl on reinforcer latency. These results also suggest that the addition of 0.32 mg/kg xylazine to 0.032 mg/kg fentanyl may alter motivation for sucrose. Interestingly, people who use drugs report that xylazine-adulterated fentanyl can be identified because it causes dryness of the mouth (Friedman et al. 2022), which is a well-known side effect of α-2 adrenergic agonists (Scully 2003). Therefore, it is also possible that reduced latency to complete an FR10 schedule for 10% sucrose solution when xylazine was added to fentanyl was driven by thirst, but further experiments would be needed to confirm this theory.

### There were no major sex differences in these experiments

Another major strength of these experiments was the use of both male and female rats because few fentanyl discrimination studies have been conducted in both males and females, and those that have reported conflicting results (Burgess et al. 2024; Schwienteck et al. 2019). In one study in rats trained to discriminate morphine, female rats were more sensitive to the interoceptive effects of morphine, but a follow-up study determined this difference was likely due to augmented suppression of response rate in males that affected learning and not to differences in the interoceptive effects of morphine (Craft et al. 1996; Craft et al. 1998b; Craft and Stratmann 1996). Additionally, the ability of fentanyl to promote morphine-like effects did not differ between males and females (Craft et al. 1996). Our results did not indicate sex differences in fentanyl discrimination or in the effects of xylazine on fentanyl discrimination. We did note that, although the proportion of male and female rats that met testing criteria after 0.01 mg/kg fentanyl training was not statistically significant, more male rats tended to meet testing criteria after 0.01 mg/kg fentanyl training, which may be worth further investigation, especially at different training doses. Therefore, future experiments could examine sex differences in rats trained on lower doses of fentanyl. Additionally, the effect of 0.1 mg/kg xylazine on the ED50 of fentanyl was more variable in males than in females, despite no difference in the mean between males and females, which suggests that there may be some individual differences in males in sensitivity to the impact of xylazine on the interoceptive effects of fentanyl. Interestingly, there is evidence for sex differences in kappa opioid receptors and their effects, including enhanced sensitivity to the depressive and interoceptive effects of kappa opioid agonists in males (Craft et al. 1998a; Rasakham and Liu-Chen 2011; Russell et al. 2014). Future experiments could reveal if this sex difference in kappa opioid effects could explain this increased variance in males or the only other major sex difference in the present study, the increased fentanyl-appropriate responding after 1 mg/kg xylazine administration in females compared to males.

### Limitations and considerations

A limitation of the present study is that we only investigated the effects of xylazine on fentanyl interoception 10 minutes after drug administration. Anecdotal reports from people who use drugs suggest that the addition of xylazine to fentanyl enhances the effects of the drug by making it feel subjectively more like heroin (Friedman et al. 2022). Notably, heroin is less potent and has longer-lasting effects than fentanyl, but the effects of heroin can substitute for fentanyl (AlKhelb et al. 2022; Fairbairn et al. 2017; Flynn and France 2021; 2022). Therefore, xylazine could extend the interoceptive effects of fentanyl across time, which remains to be tested. Another limitation is that we did not train separate groups of rats on the different training doses fentanyl. The training dose of a drug can be a major factor in which interoceptive effects of a drug are used to guide lever choice and can impact the median effective dose (Colpaert et al. 1980; Solinas et al. 2006; Stolerman et al. 2011). Interestingly, there is evidence that pretreatment with the opioid antagonist naloxone does not completely block fentanyl-appropriate responding in rats trained to discriminate lower doses of fentanyl (0.005-0.01 mg/kg), which suggests that the interoceptive effects of these lower doses may not completely rely on the direct effects of fentanyl on opioid receptors (Colpaert et al. 1980). Therefore, future experiments may evaluate the effects of xylazine on fentanyl discrimination in rats trained to discriminate a lower dose of fentanyl.

### Conclusion and clinical implications

Overall, these results suggest that xylazine potentiates sensitivity to the interoceptive effects of fentanyl. Importantly, these findings support evidence from clinical reports that suggest one reason for xylazine adulteration is to enhance the effects of fentanyl (Friedman et al. 2022). This preclinical model could be used to determine a biological mechanism for the effects of xylazine on fentanyl interoception, which is important for understanding and treating substance use disorders and overdoses involving the use of xylazine-adulterated fentanyl. Most people who use opioids consider xylazine adulteration to be unwanted (Spadaro et al. 2023; Tan et al. 2024). This is likely because too much xylazine adulteration can result in heavily sedative effects (Friedman et al. 2022). Therefore, the ratio of xylazine to fentanyl may be an important factor in its effects, but this is difficult to study in clinical populations. This preclinical model could be used to investigate differences between the interoceptive effects of varying ratios of xylazine to fentanyl. Importantly, we also show evidence that xylazine may inhibit the interoceptive and/or rewarding effects of higher doses of fentanyl, which could have important implications for risk of overdose in people using xylazine-adulterated fentanyl.

## Acknowledgements and funding

This project is supported by the Food and Drug Administration (FDA) of the U.S. Department of Health and Human Services (HHS) as part of a financial assistance award [Research Triangle Center of Excellence in Regulatory Science & Innovation, U01FD007857] totaling $1,453,116 from Center for Drug Evaluation and Research (CDER). The contents are those of the authors and do not necessarily represent the official views of, nor an endorsement, by FDA/HHS, or the U.S. Government. This project was also supported by the Bowles Center for Alcohol Studies (JB, ZAM). BNB was supported by the National Institute of General Medical Sciences K12 GM000678 (PIs: Donald T Lysle and Kathryn J Reissner). We would like to acknowledge Caroline Krieman and Alejandro Mosera for assistance with investigation.

## Author Contributions (CRediT)

**BNB**: Conceptualization, methodology, formal analysis, investigation, data curation, writing - original draft, writing - review & editing, visualization, supervision. **JMC**: Investigation, data curation. **MLB**: Conceptualization, resources. **ZAM**: Conceptualization, resources, supervision, project administration, funding acquisition. **JB**: Conceptualization, methodology, formal analysis, resources, writing - review & editing, visualization, supervision, project administration, funding acquisition.

